# Data-driven characterization of molecular phenotypes across heterogenous sample collections

**DOI:** 10.1101/248096

**Authors:** J. Mehtonen, P. Pölönen, S. Häyrynen, J. Lin, T. Liuksiala, K. Granberg, O. Lohi, V. Hautamäki, M. Nykter, M. Heinäniemi

## Abstract

The existing large gene expression data repositories hold enormous potential to elucidate disease mechanisms, characterize changes in cellular pathways, and to stratify patients based on their molecular profile. To achieve this goal, integrative resources and tools are needed that allow comparison of results across datasets and data types. We propose an intuitive approach for data-driven stratifications of molecular profiles and benchmark our methodology using the dimensional reduction algorithm t-SNE with multi-center and multi-platform data representing hematological malignancies. Our approach enables assessing the contribution of biological versus technical variation to sample clustering, direct incorporation of additional datasets to the same low dimensional representation of molecular disease subtypes, comparison of sample groups between separate t-SNE representations, or maps, and characterization of the obtained clusters based on pathway databases and additional multi-omics data. In the example application, our approach revealed differential activity of SAM-dependent DNA methylation pathway in the acute myeloid leukemia patient cluster characterized with *CEBPA* mutations that accordingly was validated to have globally elevated DNA methylation levels.

## Introduction

Gene expression profiling represents the most common genome-wide method for studying cells in healthy and disease states. As a result, large data repositories for data sharing across studies exist (1–3). However, in practice the major limitation to utilizing these results is the lack of integrative tools to compare molecular disease profiles across datasets and for including additional studies and data types into the analysis.

One common approach to discover different cellular states and disease types based on gene expression is to use unsupervised methods that require no prior knowledge on sample groups within a dataset (4). In this way, new molecular subtypes can be discovered in an unbiased manner. Dimensionality reduction belongs to such data-driven methods and is well-suited for discovery of sample grouping from complex high-dimensional gene expression data. Currently, there are several computational methods available for this task (reviewed in 5). Given a high-dimensional input matrix containing several thousand gene expression values, the algorithms use measures of (dis)similarity and an optimization strategy to return sample coordinates in a lower dimension, in a manner that optimally preserves their relative placement in the original coordinate space. To be useful, data points appearing similar (proximate) in the lower dimensional visualization should be trusted to be similar in actuality. In addition, all original proximities ideally would become visualized close-by. These properties can be captured using the metrics of trustworthiness and continuity (6), analogous to precision and recall in classification. However, in context of data generated by multiple laboratories, the contribution of technical variation to the obtained visualization remains a challenge (7).

We propose here an approach that incorporates data on lower dimensional representations obtained with the well-performing dimensionality reduction algorithm t-stochastic neighbor embedding (t-SNE) (8) and benchmark the methodology with multi-center and multi-platform data representing hematological malignancies. Our framework enables choice of analysis parameters to minimize batch effects that arise from different laboratory protocols, incorporation of additional samples to previously defined t-SNE space and integration of data from different measurement platforms.

## Methods

### Datasets

A dataset of 9,544 gene expression profiles from the Gene Expression Omnibus (GEO) database (1), comprising patient samples representing different cancers and proliferative disorders of hematopoietic lineage origin, cell lines and normal blood cell types represents the main data used for method testing. We refer to this sample set as Hemap in the following text. The curated sample annotations and disease categories are available at http://compbio.uta.fi/hemap/. These data represent microarray data from the commonly used hgu133Plus2 platform that were processed using the RMA probe summarization algorithm (9) with probe mapping to Entrez Gene IDs (from BrainArray version 18.0.0, ENTREZG) to generate gene expression signal levels. Next, a bias-correction method (10) developed for clinical microarray data was applied to correct for technical differences between studies. Additional 98 biological replicate samples from these studies, and all 108 samples from the study GSE49032 (11) were left out as validation data. A microarray dataset (12) from a different array platform (hgu133a+b) and RNA-seq data from the TCGA AML study (13) were included to benchmark robustness in discovery of molecular subtypes and addition of samples from different measurement platforms.

### Inclusion of new data from the same measurement platform

To benchmark integration of new samples from the same microarray platform, the left-out samples were used (refer to **Table S1**). The data was normalized as similar as possible to that applied to the original data. Ideally, RMA summarization (9) and bias correction (10) of new samples should be performed together with the original samples. However, considering the dimensions of the original data (9,544 samples), co-normalization of new and old samples is not convenient from the viewpoint of memory usage and computational complexity. In addition, regeneration of the full data matrix would also require re-running all the downstream analysis to maintain the consistency of the results. Rather, we revised the preprocessing approach that allows normalization of new samples to the space of original data. The background correction step of RMA was performed in a standard way as it requires no inter-sample information. The quantile-normalization step, however, utilizes information across all samples. In normalizing novel samples, we used the normalized distribution from the original data to ensure that data distributions for novel samples do not differ from those of the original ones. In the median polish summarization, likewise, the row (probe) effect of the original data is used instead of calculating it across the novel samples. Otherwise median polishing is performed as usual. In the bias correction step of novel samples, the coefficients describing the dependency between the bias metrics and gene expressions were obtained from the original data set. It should be noted that all the samples to be normalized should also meet the quality control requirements that were used with the original data set.

### Inclusion of new data from different measurement platforms

The TCGA RNA-seq data (13) was obtained through cghub and realigned to hg19 genome using Tophat2 (14) version 2.0.12 with default parameters. The expression of genes included in the microarray was calculated by counting the reads aligning to the corresponding probe regions. RNA-seq data were further processed by log2 transformation. Data from hgu133a and b platform was normalized using RMA with probe mapping to Entrez gene IDs as above.

### Quantitative metrics for assessing dimensionality reduction and clustering results

To account for both biological and technical differences that are characteristic of sample sets generated by different studies, we propose two new metrics to guide the feature selection for dimensionality reduction in context of such heterogeneous data:

*NMIp*: This metric was calculated to assess how well the obtained sample clustering can distinguish known biological subtypes (phenotypes) based on normalized mutual information (15) (NMI) between cluster assignment and class labels (maximized).

*NMIe*: Similar as above, NMI was calculated between cluster and experiment (data series) identifiers. The data series differences represent mainly technical (not biological) differences between samples and therefore this measure was minimized.

Initial comparison of the different dimensionality reduction methods encouraged the selection of t-SNE method, specifically the Barnes-Hut approximated version of t-SNE implementation (BH-SNE) (16), to serve as a benchmark scenario for data-driven exploration of disease subtypes. We compared the default step that uses Principal Component Analysis (PCA) (17) for initial reduction of features to selection of genes based on variance. Selecting of 20 to 100 PCs or 2.5 to 50% of the most variable genes were compared.

Kernel density-based clustering algorithm known as mean-shift clustering (18) with bandwidth parameter set to 1.5 (subsets of data, one cancer type) or 2.5 (all data) was used (LPCM-package in R) to cluster the data following the dimensionality reduction. This method allows the discovery of sample sets which share similar features without having to pre-specify the number of clusters. The term “cluster” is used in the text to refer to this computational clustering result, and the term “group” is used in context of visual examination.

### Remapping of samples to the t-SNE maps

New samples were mapped to an existing t-SNE space by using a modified version of the BH-SNE (16) implementation. While the original algorithm initializes the embedding points by sampling from a Gaussian distribution, our implementation for adding new samples used the established map for initializing the embeddings for existing samples. Embeddings of existing samples were kept locked throughout the run of the gradient-descent optimizer, while the embeddings for new samples were computed in parallel and independently of each other. Independent embeddings of new samples do not affect each other or the embeddings of locked samples, thus preserving the structure of the established t-SNE map.

The t-SNE method minimizes the divergence between two distributions: a distribution that measures pairwise similarities between the original data objects (by default Euclidean distance is used) and a distribution that measures pairwise similarities between the corresponding points in the embedding. To remap RNA-seq samples, Euclidean distance was replaced by correlation as a distance measure (one minus Pearson correlation between samples). By using a distance metric based on correlation, the similarities between RNA-seq samples and different microarray samples could be estimated without further transformations or normalizations of RNA-seq data.

### Gene set analysis

The gene lists for the characterization of sample clusters were obtained from MsigDB v5.0 (19), Wikipathways (06.2015) (20), Recon 1 (21), Pathway Commons 7 (22) and DSigDB v1.0 (23). Gene sets were limited to contain between 5 to 500 expressed genes per gene set, resulting in 19,680 gene sets that were evaluated across the dataset. In addition, gene sets were defined on basis of significant cluster correlation. The gene set variation analysis (GSVA) (24), available in the R/Bioconductor package GSVA 1.13.0, was used to assign a gene set enrichment score (positive for increased and negative for decreased expression) in a sample-wise manner with the following settings: mx.diff=F, tau=0.25, rnaseq=T if RNA-seq, otherwise rnaseq=F. Empirical *P*-value was computed using 1000 random permutation of genes within the gene set. Estimation of significance was limited to a range of gene set sizes (5-20, 25, 30, 40, 50, 75, 100, 200, 300, 400, 500) to adequately account for differences in gene set size distribution. The observed pathway score was compared with the random permutations of a corresponding gene set size and empirical *P*-value computed as the number of higher/lower scores in the permuted set divided by the total number of permutations. Enrichment of significant scores in a specific cluster was computed using a hypergeometric test.

### Correspondence between t-SNE-map clusters

Similarity in sample clustering between t-SNE maps was evaluated in a data-driven manner using GSVA (24) enrichment scores. Two gene sets (20 top ranked positively or negatively correlated genes, separately) were defined for each t-SNE map cluster based on significant Pearson correlation to assess the robustness of the clustering and correspondence between the maps. Next, enrichment of other gene sets was compared: a nominal *P*-value cutoff 0.05 was used for the single data series with sample sizes less than 100. Next, these gene sets were evaluated from the Hemap dataset, requiring the same directionality in the correlation (to corresponding cluster) and adjusted *P*-values less than 0.001.

### Characterization of sample clusters based on multi-omics features

Categorical or binary annotation features (including clinical variables), continuous/discrete numeric expression values (14,274 genes), continuous/discrete numeric molecular data (mutation, CNV, methylation and chromosomal translocation) values or continuous GSVA pathway scores (19,680 pathways) were collected for each sample. Missing values were marked as NA. Each t-SNE map cluster (binary feature) was tested against all other features and the statistical significance evaluated using Spearman correlation between a binary cluster feature and a numeric or binary other feature. Multiple hypothesis testing correction was performed using the Bonferroni method, using the total number of comparisons *N* × *M*, where *N* is the number of binary cluster features tested and *M* is the number of features within the feature type tested. Pairwise analysis results were filtered using the adjusted *P*-value cutoff 0.001 for Hemap data and 0.05 for the smaller TCGA AML dataset (13). For gene sets, the cluster to gene set enrichment test was used as an additional filter as described before.

### Kaplan-Meier survival analysis

Survival time and status for each TCGA AML sample was obtained from the supplementary table from the original publication (13) (“Patient Clinical Data” dated 3.31.12). The R package ‘survival’ was used to compute Univariate Kaplan Meier curves for each TCGA cancer-map cluster and to calculate the log-rank test.

### Discretizing methylation signal levels with mixture models

Gaussian finite mixture models were fitted by expectation-maximization algorithm provided in the R package mclust (25) (version 4.3) to identify whether the value obtained for a given methylation probe belonged to the signal (expressed) or noise distribution in each sample. The model was chosen by fitting both equal and variable variance models and ultimately choosing the model which achieved a higher Bayesian Information Criterion (BIC) to avoid overfitting. A model with three components was fitted by choosing either an equal or unequal variance model according to BIC. After model fitting, the percentages of measurements belonging to each component were calculated for each sample. The portions of highly methylated regions in the studied cluster were compared against the rest of the samples by calculating the Mann-Whitney U test.

### Code availability

All custom code for analysis and normalized data matrices (RData files) for reproducing the analysis presented will be freely accessible upon publication of the manuscript.

## Results

### Assessing the contribution of technical variation in sample separation

A fundamental challenge for joint analysis of the genome-wide data available in public repositories is the technical variability between data generated by different laboratories. To develop robust data integrative solutions, we use here a microarray dataset, Hemap, that was collected across several studies for researching disease mechanisms in context of hematological malignancies (see **Methods**). The data was jointly processed, including established normalization (9) and batch effect correction (10) steps. To organize the disease subtypes in a data-driven manner, we used the dimensionality reduction method t-SNE that has been successfully applied in context of both simulated and real datasets (8). The challenge in application of t-SNE across the heterogeneous data is illustrated with the subset of Hemap AML samples upon iterative addition of experiments in **Fig. 1a**. A PCA pre-processing step is included to the default implementation, which in this case results in separation of the data based on the study (**Fig. 1A**, PCA BH-SNE). Testing alternative solutions, we found that selection of genes based on variable expression results in sample grouping that does not reflect data origin (**Fig. 1A**, Variable genes BH-SNE). The technical bias in the PCA BH-SNE result, was not apparent based on two common metrics, continuity and trustworthiness (6) (PCA BH-SNE 0.92 and 0.96; Variable genes BH-SNE 0.95, 0.97, respectively). To address this issue, we defined two new quality measures for quantitative evaluation of the sample clustering in context of heterogenous datasets: NMIp that captures the separation of phenotypes (maximized), and NMIe that can be used to penalize the separation of data by experiment (minimized) (see Methods). The calculation of these metrics requires that at least some samples have an annotated class and that origin (study/experiment identifier) of the samples is known. The effect of choosing different number of principal components, or different percentage of genes, is compared using these metrics in **Figure 1B** for the AML subset and the full sample collection. In both cases, feature selection based on genes with highest variance performed favorably, reducing the technical biases. Therefore, we selected 15% most variable genes for t-SNE analysis in this study. Overall, the obtained AML t-SNE map separated clinical subtypes in 38/38 (100%) of cases in an unsupervised manner, similarly to the supervised classification with a Prediction Analysis for Microarrays (PAM) classifier (26) (**Table S2**). The annotated sample category is visualized in **Figures 1C-E** on t-SNE maps, including two other sample subsets (acute lymphoblastic leukemia (ALL) and lymphomas). In each, a sample grouping driven by cancer subtypes was obtained. Therefore, the quantitative assessment based on the new metrics can guide parameter selection for unsupervised sample stratification methods to mitigate technical variation effects.

**Figure 1.**
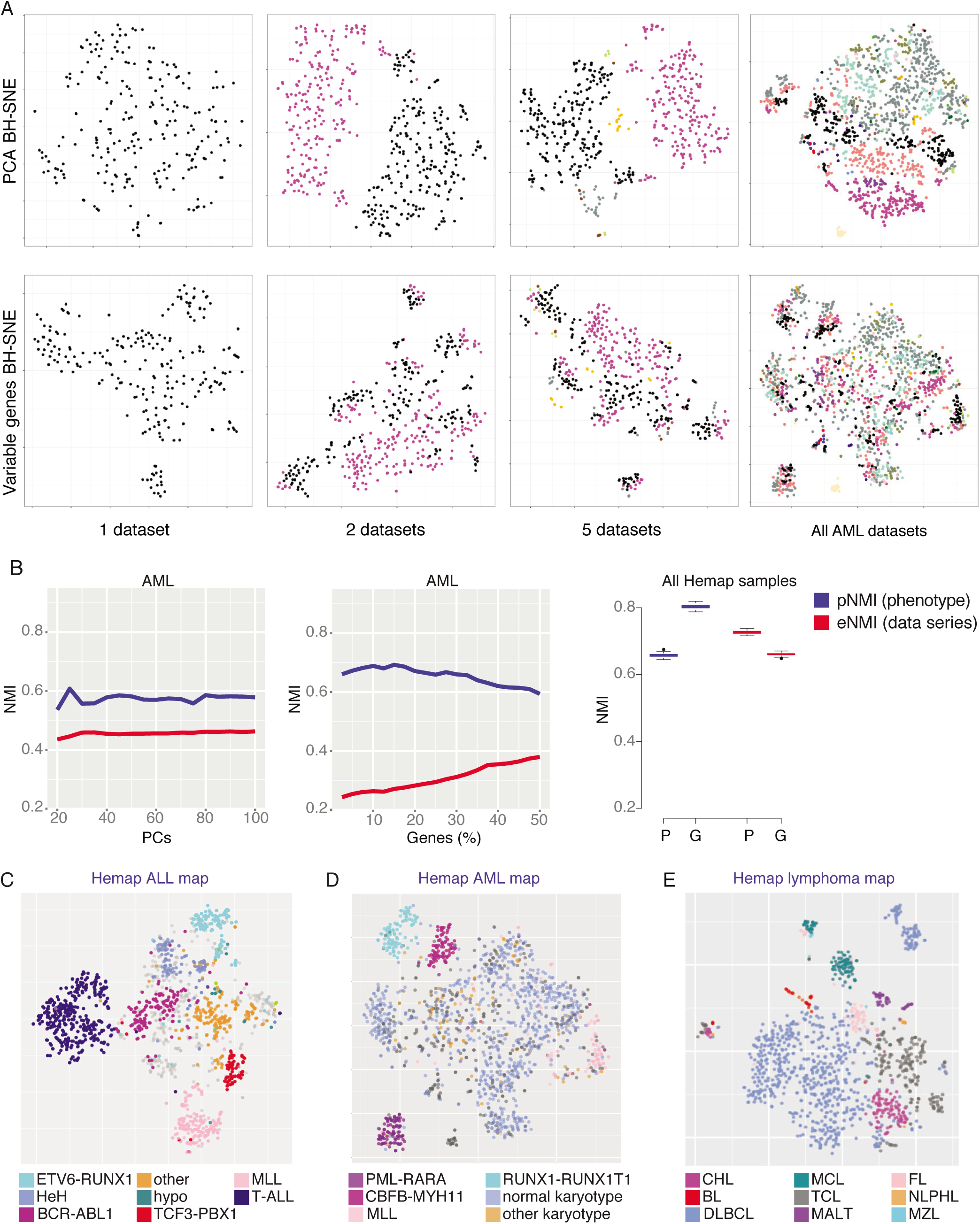
Assessing the contribution of technical variation in sample clustering. **A**. Iterative addition of AML data series to the sample set used for t-SNE is shown. The difference between using the PCA preprocessing step (above) or using 15% of most variable genes (below) is visualized from a succession of maps (a single data series, 2, 5 or all). **B**. The metrics for separation of phenotypes (NMIp, in blue) and data series (NMIe, in red) shown represent the mean of 30 permutations with different seed selections. Selecting a different number of principal components (PCs) is compared to selecting a different percentage of variable genes using the Hemap AML sample set. For all Hemap samples, BH-SNE results using PCA (P) or 15% most variable genes (G) are compared. The fine structure on the t-SNE maps with 15% gene selection matches closely pre-B-ALL (**C**), AML (**D**) and BCL (**E**) clinical subtypes (in color).

### Incorporation of additional datasets to predetermined t-SNE space

We next asked whether the clustering of disease (sub-)types on the t-SNE maps could be used to characterize new samples. We developed a remapping algorithm that allows additional samples to be included to the existing map (see Methods). The left-out samples (*N*=10 from AML) were all assigned to the same cluster as their replicates on the Hemap AML t-SNE map (**Fig. 2A**, diamond shapes, see also **Table S3** and **Fig. S1** that shows the successful association of all 98 validation samples with the disease-of-origin). Re-mapping an independent microarray dataset to the ALL t-SNE-map successfully assigned the clinical subtype for 95% of samples (**Fig. 2B**). Since many new studies currently use RNA-seq, the remapping algorithm was extended to incorporate alternative types of data by revising the similarity metric (see Methods). From AML RNA-seq samples (13), 88% were placed to a cluster matching the annotated category (**Fig. 2C**). In this manner, our approach readily extends beyond the present sample set and shows robustness for different technologies.

**Figure 2.**
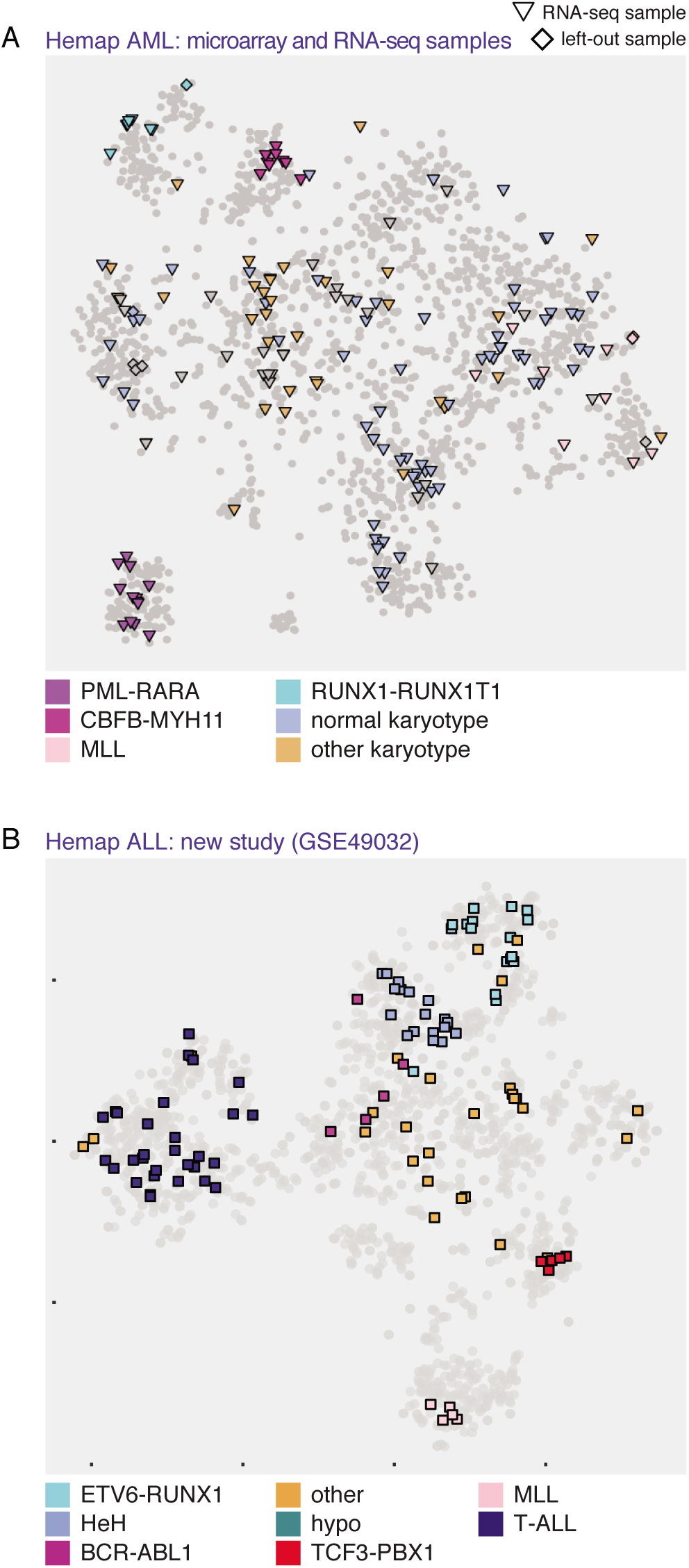
Remapping new samples to the t-SNE map. **A**. AML microarray validation samples that were left out (*N*=10, diamonds) and TCGA RNA-seq samples^13^ (*N*=162, triangles) re-mapped on the Hemap AML t-SNE map. Notice that similar samples mapped in close proximity to each other may overlap on the visualization. **B**. Remapping result for samples from an independent ALL study^11^ (GSE49032) to the Hemap ALL t-SNE map. The subtype of re-mapped samples is indicated in color in **A** and **B**.

### Comparison of data-driven stratifications of molecular disease subtypes

Comparisons between studies to determine whether the same molecular subtypes segregate in a reproducible manner was considered next. Towards this end, we developed methodology for examining the correspondence between clusters on separate t-SNE maps. We chose the TCGA AML (13) (RNA-seq, **Fig. 3**) and Ross ALL (12) (microarray, **Fig. S2**) studies for comparative analysis. **Figure 3A** shows the generated t-SNE map for TCGA RNA-seq samples, which resulted in seven distinct clusters (referred to as TCGA clusters 1-7). The comparison with annotated cytogenetic types was consistent; samples carrying PML-RARA, RUNX1-RUNX1T1, CBFB-MYH or MLL fusions segregated into distinct clusters. To identify matching clusters from the Hemap AML t-SNE map, we developed a data-driven approach that defines and scores gene sets for each cluster (see Methods, refer to **Table S4** for TCGA cluster gene sets). The samples with significant enrichment are colored on the HEMAP AML t-SNE map, and their enrichment score compared to annotations as an oncoprint (**Fig. 3B**). Samples with matching fusion gene status received the highest enrichment scores, allowing matching of clusters between the maps (**Table S5**). The application to ALL samples (12) (**Fig. S2**) showed similar robustness in discovery of matching subtypes.

**Figure 3.**
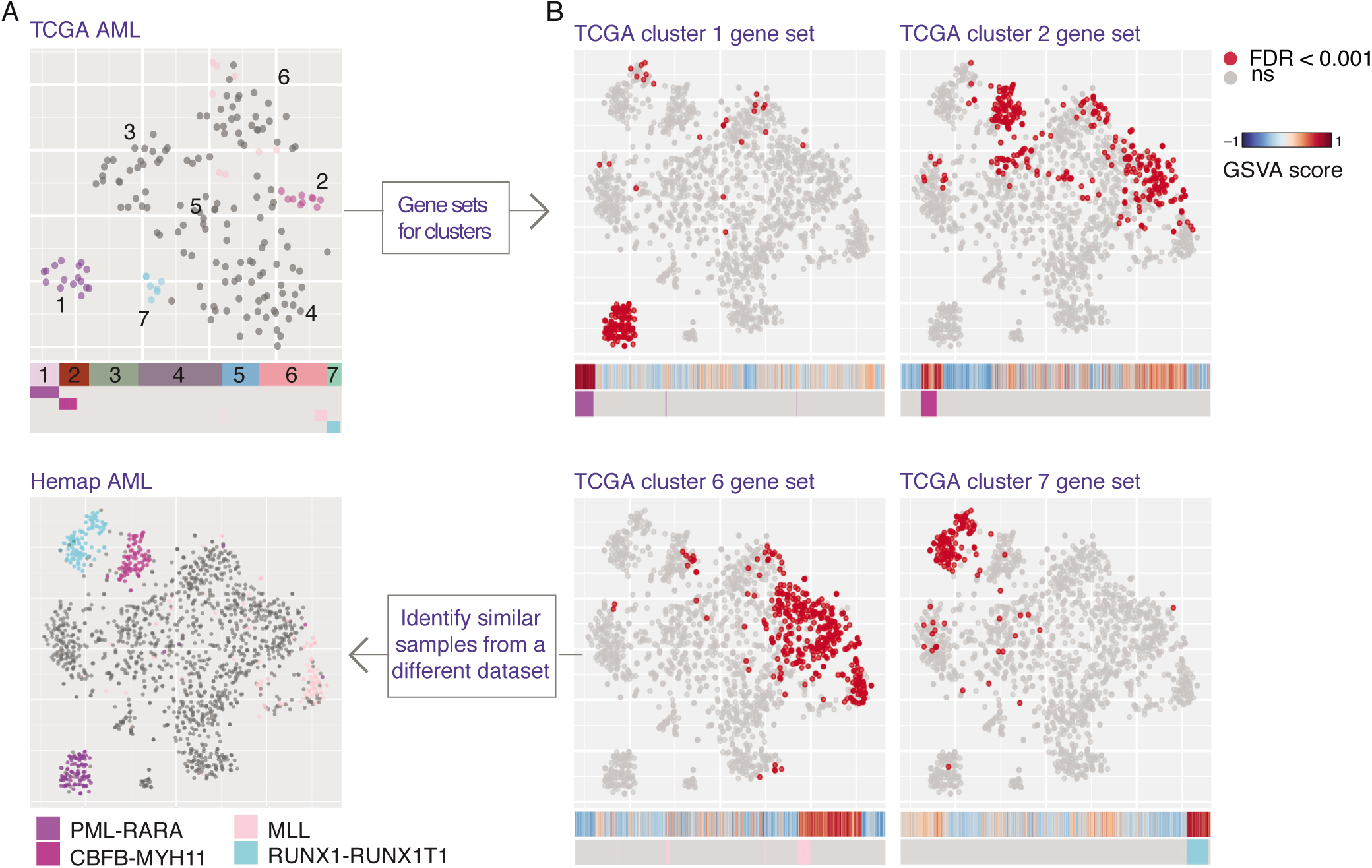
Evaluation of disease subtypes across datasets. Comparison of sample clustering on the t-SNE maps for TCGA AML RNA-seq samples (*N*=162) and Hemap AML is shown. **A**. The data-driven cluster assignment (TCGA clusters 1-7) can be compared with sample molecular annotations colored on the map and the oncoprint heatmaps below. **B**. Enrichment scores for TCGA cluster gene sets that matched samples with common fusion genes (clusters 1, 2, 6 and 7) are colored on the Hemap AML map (significant enrichment adj. P-value < 0.001 in red). The raw GSVA scores are shown below as an oncoprint heat map (red tones indicate high expression of the gene set and blue tones low expression).

### Integration with multi-omics profiles and pathway activity analysis to characterize the discovered molecular subtypes

The clinical classification of AML has traditionally distinguished between fusion gene-positive categories (27). However, additional clusters segregate in the TCGA and Hemap AML t-SNE maps (**Figs. 3A** and **C**) that can be matched between the studies (**Fig. 4A, Table S5**). Statistically significant associations between the map clusters and different multi-omics features or pathway activity scores were queried to characterize the discovered patient groups (see Methods). Based on correlation of cluster assignment with multi-omics TCGA data (13), *NPM1* mutations were associated with two clusters (TCGA clusters 3 and 6), whereas *CEBPA* mutations characterized TCGA cluster 5 (**Fig. 4B**). This could be confirmed in corresponding Hemap AML clusters (**Fig. 4C**). The distinction between the two NPM1 positive clusters was associated with the cellular morphology (FAB type in **Fig. 4B**). In addition, our analysis revealed a subgroup (TCGA cluster 4 and corresponding Hemap samples indicated in **Fig. 4A**) with several patients positive for either *TP53* or *RUNX1* mutations and/or complex karyotypes (**Figs. 4B**-**D**). The distinction between these non-fusion samples is clinically relevant, as the identified clusters differed in overall survival (**Fig. 4E**).

**Figure 4.**
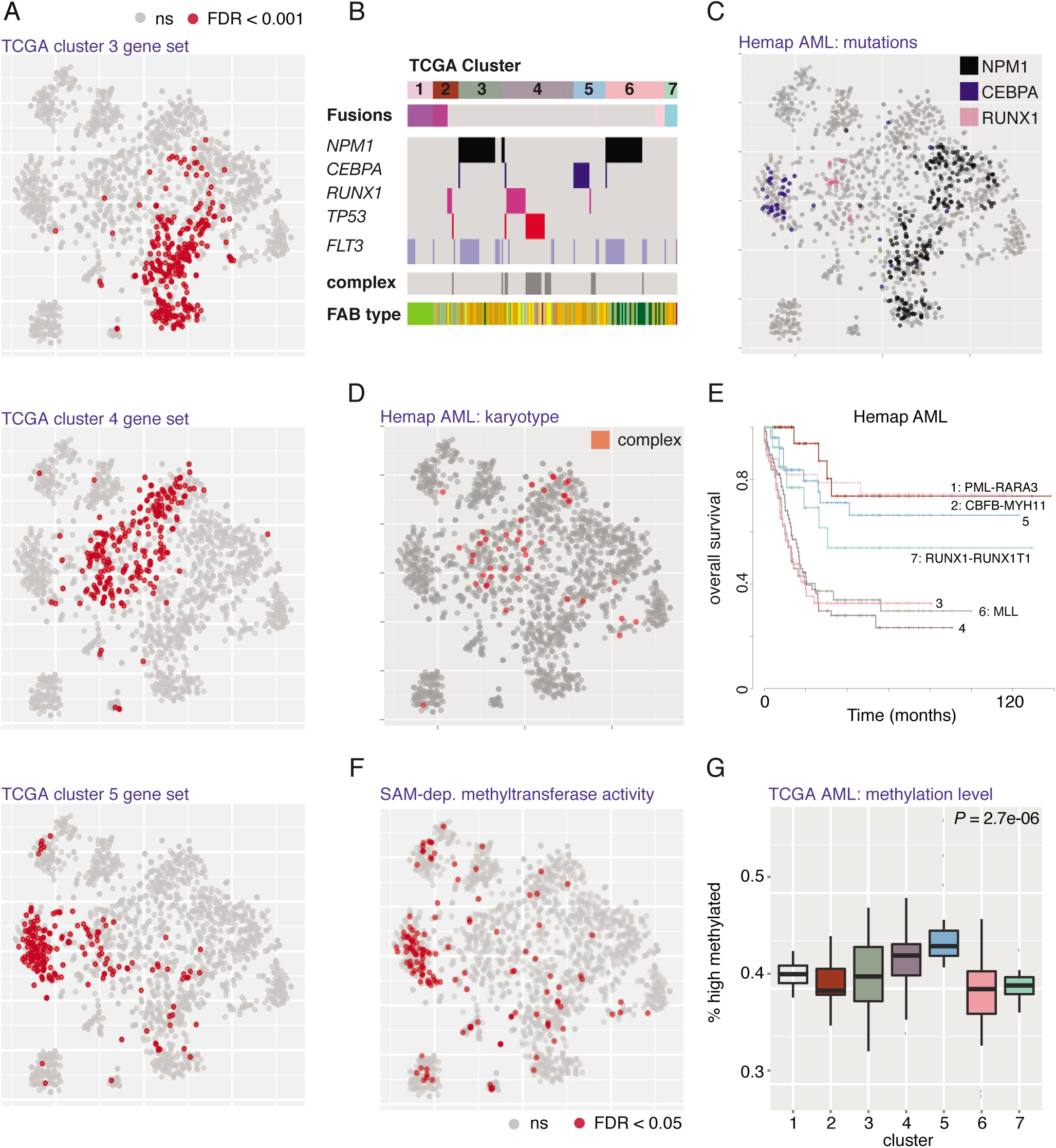
Multilevel data integration reveals AML clusters with distinct mutation and epigenetic phenotypes. **A.** Significant enrichment (adj. P-value &0.001) of TCGA clusters 3-5 gene sets is colored on the Hemap AML map (as in Fig. 3B). **B**. Oncoprint heatmap comparing fusion gene, mutation, karyotype and phenotype (FAB type) of samples in different TCGA clusters. **C**. Mutational status for most significantly cluster-associated mutations is indicated in color on the Hemap AML map. **D**. Location of complex karyotype samples indicated in color on the Hemap AML t-SNE map. **E**. The survival analysis of Hemap patients comparing the clusters matched with the TCGA map are shown as Kaplan-Meyer plots. **F**. Significant enrichment (adj. P-value < 0.05) for SAM-dependent methyltransferase activity is shown as in **A**. **G.** DNA methylation levels compared between TCGA AML clusters. The percentage of regions assigned to the high methylation state compared between TCGA cluster 5 patients (*N* = 20) and rest of TCGA AML clusters (Mann-Whitney U-test *P*-value is indicated).

Next, we detected significant associations between the cluster category and pathway activity. The gene set enrichment analysis (see Methods) confirmed that the TCGA and Hemap patients from the matched clusters share interesting molecular characteristics (**Table S5**). Intriguingly, TCGA cluster 5 (*CEBPA* mutated, **Fig. 4B**) and its corresponding cluster on the Hemap AML map (refer to **Figs. 4A** and **C**) were significantly associated with S-adenosylmethionine (SAM)-dependent methyltransferase activity (**Fig. 4F,** hypergeometric test adjusted *P*-values 5.6e-05 and 1.5e-26, respectively). Because all DNA methyltransferases use SAM, we quantified TCGA DNA methylation data to test whether global changes in methylation patterns exist between clusters (see **Methods**). Accordingly, we observed a significant elevation in DNA methylation level in the TCGA cluster 5 compared to other samples (**Fig. 5G**, Mann-Whitney U-test *P*=2.681e-06).

## Discussion

A large number of biological conditions have been characterized at genome-wide level since the introduction of microarray and deep sequencing technology (1–3), However, most of the studies include only tens to hundreds of samples. Cancers of hematopoietic origin serve as an important example where data integration is essential from the sample availability perspective, as many of these cancers are rare on the population level (27,28). Thus, understanding the complete heterogeneity and similarity of diseases states and their subtypes requires integrative data analysis methodology. Here, we tested solutions that allow distinguishing technical variation and evaluating the robustness of the obtained biological stratifications between studies, comparative analysis of pre-existing patient molecular data and inclusion of new sample sets and data types, as they become available.

Removing technical variation, i.e. batch effects, from data before downstream analysis is a vital part of studies analyzing multiple datasets generated from different experiments (7,10,29). We propose here two NMI metrics that can guide the choice among alternative methods that achieve minimal loss of biologically relevant gene information (high separation of phenotypes) while at the same time reduce the technical variation enough not to interfere with the biological interpretation of the results. In the analysis presented, after data normalization and batch correction, we further remedy the technical biases by removing less variable genes as a preprocessing step for t-SNE (8). Evaluating the dimensionality reduction alone (continuity, trustworthiness (6)) was found insufficient to capture the technical bias that persisted when the default PCA pre-processing step was used. Alternatively, decomposing the variation could be attempted to reduce batch effects, including methods that attempt to remove the variation contained in top principal components (30,31). Our approach to evaluate the obtained sample clustering does not make any assumptions about the data. Therefore, it extends to more than just microarray and RNA-seq data presented here. For example, it will be very relevant in single-cell context where the challenge of batch effects has been recognized (32) and new data sets are becoming increasingly available.

To date, microarray repositories are still by far the largest resource for molecular data and hold a vast potential for large-scale studies. Compared to existing transcriptome collections (33,34), our goal was to develop methods that can generalize the integrative analysis beyond the initial dataset. We showed that in the context of the popular dimensionality reduction method t-SNE, new samples can be added to extend the existing map. Moreover, as next-generation sequencing based assays are gaining ground, we also present methodology to include RNA-seq profiles for joint analysis with microarray results. Secondly, we introduce a gene set-based method to compare clusters between different datasets, using the unsupervised t-SNE projection and clustering as the initial starting point, and identifying corresponding clusters using gene set enrichment. In both cases, there are considerations related to how representative the initial dataset (for remapping or defining cluster gene sets) are of the biological context, such as the cancer subtypes considered here. The enrichment results can be different if the sample composition changes, as genes are ranked based on kernel estimation of the cumulative density function using all samples (24). For comparing samples from small studies to a larger reference study, the re-mapping approach would be better suited.

Finally, we demonstrate the integration of additional data types to further characterize the identified sample groups. Our approach considers gene expression levels as the main phenotype that gene set and pathway analysis can further characterize. Other data types (binary and continuous features such as mutations and methylation levels) were included by correlating them with the cluster observed in this phenotype (gene expression) space. Alternatively, many multi-view methods have been introduced that are designed to preserve structures between several layers of data (35,36). While these methods appear theoretically promising, in practice different data types and result interpretation present challenges (36). Our analysis in AML revealed new insight on molecular subtypes, revealing a distinct clustering of *CEBPA*, *NPM1*, *RUNX1* and *TP53* mutation positive samples. These molecular phenotypes that we found to distinguish fusion gene negative samples matched a distinct gene expression state. The gene set scoring allowed the same patient groups to be identified from the Hemap sample AML map, further demonstrating that the molecular subtypes found are robust, in a comparable manner to the fusion genes distinguished in previous classifications (26,27). The cluster with the worst survival in our analysis included samples that were characterized by complex karyotypes and *TP53* mutations, agreeing with recent genotyping data of recurrent mutations (37). Furthermore, the analysis of cluster-associated mutations distinguished a patient cluster with *CEBPA* mutations. Using pathway analysis, we could demonstrate that TCGA and Hemap patients from the matched clusters had elevated expression of genes involved in SAM-dependent methylation activity. Previous studies have found contradicting results, reporting both specific (13) and broad (38) methylation changes. We validated that patients with high pathway activity score had an elevated global DNA methylation level using multi-omics TCGA data.

In conclusion, we present new data integration approaches for multi-center and multi-omics datasets that allow researching disease subgroups. This analysis framework can be adopted to support the utilization of genome-wide data across different biological systems and disease contexts.

## Supplementary material

The supplementary material consists of the description of methods (**Online Methods**) and Supplementary display items (two figures and five tables) with their legends.

## Author contributions

M.H. and M.N. initiated the study. J.M., V.H. and M.H. designed comparison of dimensionality reduction results and J. M. performed the data analysis, P.P. performed gene set enrichment and TCGA data analysis, S.H. developed the t-SNE remapping and performed RNA-seq preprocessing and T.L. the classifier analysis. O.L. contributed to the design and analysis of AML data. M.H., M.N., and V.H. supervised the work. M.H. and M.N. wrote the manuscript. All authors commented on the manuscript.

## Acknowledgements

We would like to thank Dr. Sheila Reynolds and Roger Kramer for assistance with data analysis. The work was supported by grants from the Academy of Finland (project no. 269474 (MN), no. 276634 (MH)), The Finnish Cultural Foundation (Interdisciplinary Science Workshops, MH), Sigrid Juselius Foundation (MN), and Cancer Society of Finland (MN, MH), Nokia Foundation (VH) and University of Eastern Finland (MH).

The results published here are in part based upon data generated by The Cancer Genome Atlas project (dbGaP Study Accession: phs000178.v8.p7) established by the NCI and NHGRI. Information about TCGA and the investigators and institutions who constitute the TCGA research network can be found at http://cancergenome.nih.gov.

